# Ecology of metagenomes: incorporating genotype-to-phenotype maps into ecological models

**DOI:** 10.64898/2026.04.07.717079

**Authors:** Siqi Liu, Pankaj Mehta

## Abstract

A major theoretical problem in community ecology is to understand how genes, organisms, and environments combine to shape the structure and diversity of ecological communities. However, most classic ecological models work entirely with phenotypic parameters, neglecting the central role played by genes. This limitation is particularly acute in microbial ecology, where the widespread use of sequencing technologies allows researchers to directly measure the genomic and metagenomic properties of communities. Here, we bridge this gap by incorporating genotype-to-phenotype maps into classical ecological models, including the generalized Lotka-Volterra model (GLV) and consumer resource models (CRMs). We focus on the case where genotype-to-phenotype maps are linear, which provides a tractable yet powerful framework for analyzing complex traits. Even in this simple setting, the resulting ecological dynamics give rise to novel gene-level ecological dynamics that can be recast entirely in terms of genes, allowing us to develop an ecology of metagenomes. We find that ecological interactions between genes lead to pervasive “metagenomic hitchhiking” — low-fitness genes can survive in the ecosystem because they are integrated into genomes of high-fitness species. We also show that phylogenetic relationships between species mold the ability of closely related strains to stably coexist in complex communities. This highlights how lineage structure and competitive interactions jointly shape community composition. Our framework provides a principled foundation for interpreting metagenomic data through the lens of ecological theory.

**Author summary:** Recent advances in sequencing technologies have transformed our ability to characterize microbial communities at the genomic level. However, most classic ecological models work entirely with phenotypic parameters, neglecting the central role played by genes. Here, we address this gap by extending classical ecological models to explicitly include genotype-to-phenotype maps. We focus on complex traits where the genotype-to-phenotype map is approximately linear. We show that the resulting ecological dynamics that can be recast entirely in terms of genes, allowing us to develop an ecology of metagenomes. Our framework provides a novel perspective for interpreting metagenomic data through the lens of ecological theory.

## Introduction

Understanding the relationship between genetic composition and ecological function is one of the central challenges of modern microbial ecology [1, 2]. Genes define the identity of organisms and shape their phenotypic properties. At the same time, genes themselves are subject to ecological and evolutionary forces: they spread, persist, or vanish depending on how they influence organismal fitness and community interactions [3].

Recent advances in sequencing technologies have transformed our ability to characterize microbial communities at the genomic level. Single-cell genomics allows reconstruction of individual genomes (SAGs) from uncultured cells [4], while shotgun metagenomic sequencing profiles the entire DNA content of a community, enabling the recovery of metagenome-assembled genomes (MAGs) and the quantification of gene abundances across environments [5]. These technological advances have made it possible to observe the genetic foundations of ecology directly, highlighting a growing mismatch between data richness and the gene-free assumptions of classical ecological theory.

Classical ecological models, such as the generalized Lotka–Volterra (GLV) equations [6–8] and consumer resource models (CRM) [**?**, 9–11], have long provided powerful descriptions of species interactions and coexistence. In these models, species are treated as phenotypically distinct entities that grow and interact according to prescribed phenomenological parameters. Despite their tremendous utility, these models are fundamentally gene-free: ecological parameters such as growth rates, interaction coefficients, and resource preferences are treated as intrinsic species properties, with no connection to underlying genotypes.

These assumptions make it difficult to relate model predictions to experimental observables based on sequencing. Classic ecological models also assume the existence of well-defined species boundaries – a step that depends on taxonomic classification and becomes particularly fraught in microbial systems, where horizontal gene transfer blurs the very notion of species [12, 13]. To overcome these limitations, here we develop a theoretical framework that explicitly incorporates genomes into classical ecological models.

Building on the GLV and CRM formalisms, we represent each species by a genome vector, whose entries indicate the presence of genes. Phenotypic parameters in the ecological models are then determined by a genotype-to-phenotype (G→P) map, which translates genetic composition into ecological traits such as growth rates, resource preferences, and interaction strengths. We focus on the case where the G→P map is linear – a good approximation for complex phenotypic traits governed by many genes with small, independent effects [14, 15]. Even in this analytically tractable setting, we find that the ecological dynamics possess a surprisingly rich structure.

Within this framework, ecological interactions can be reformulated in terms of genes rather than species, giving rise to what we term an *ecology of metagenomes*. This perspective reveals several novel phenomena. In particular, we show that ecological interactions among genes lead to pervasive *metagenomic hitchhiking*, whereby low-fitness genes persist in the community by co-occurring with high-fitness genomes. We also demonstrate that multiple strains of the same species can stably coexist, offering a theoretical explanation for experimentally observed microbial diversity. We further show that the number of coexisting species is set by the rank of the G→P map, directly linking ecological diversity to the dimensionality of functional genetic space. Together, these results suggest that embedding genotype-to-phenotype maps within ecological models provides a powerful starting point towards developing a unified theory of ecological dynamics that bridges genes, phenotypic traits, and the environment.

## Models

In this section, we develop a mathematical framework incorporating linear genotype-to-phenotype maps into two classic models from theoretical ecology: the MacArthur consumer resource model (MCRM) and the generalized Lotka-Volterra (GLV) model. The resulting ecological dynamics can be formulated directly in the space of genes, allowing for the development of an ecology of metagenomes.

### MCRM with linear G→P maps

We begin by introducing the classic MacArthur consumer-resource model (MCRM), which describes how species populations grow by consuming shared environmental resources. The MCRM is one of the foundational models of ecology and forms the basis of many contemporary ecological frameworks such as contemporary niche theory [**?**] and Tilman’s resource competition theory [16].

In the MCRM, *n*_*S*_ species with abundances *N*_*i*_, where *i* = 1, 2, …, *n*_*S*_ can consume *n*_*R*_ distinct resources with abundances *R*_*α*_, where *α* = 1, 2, …, *n*_*R*_. The ecological dynamics are governed by coupled ordinary differential equations of the form

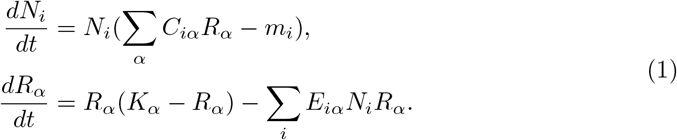

Within the MCRM, each species is defined by a set of phenotypic parameters (see Fig 1A):

**Fig 1.**
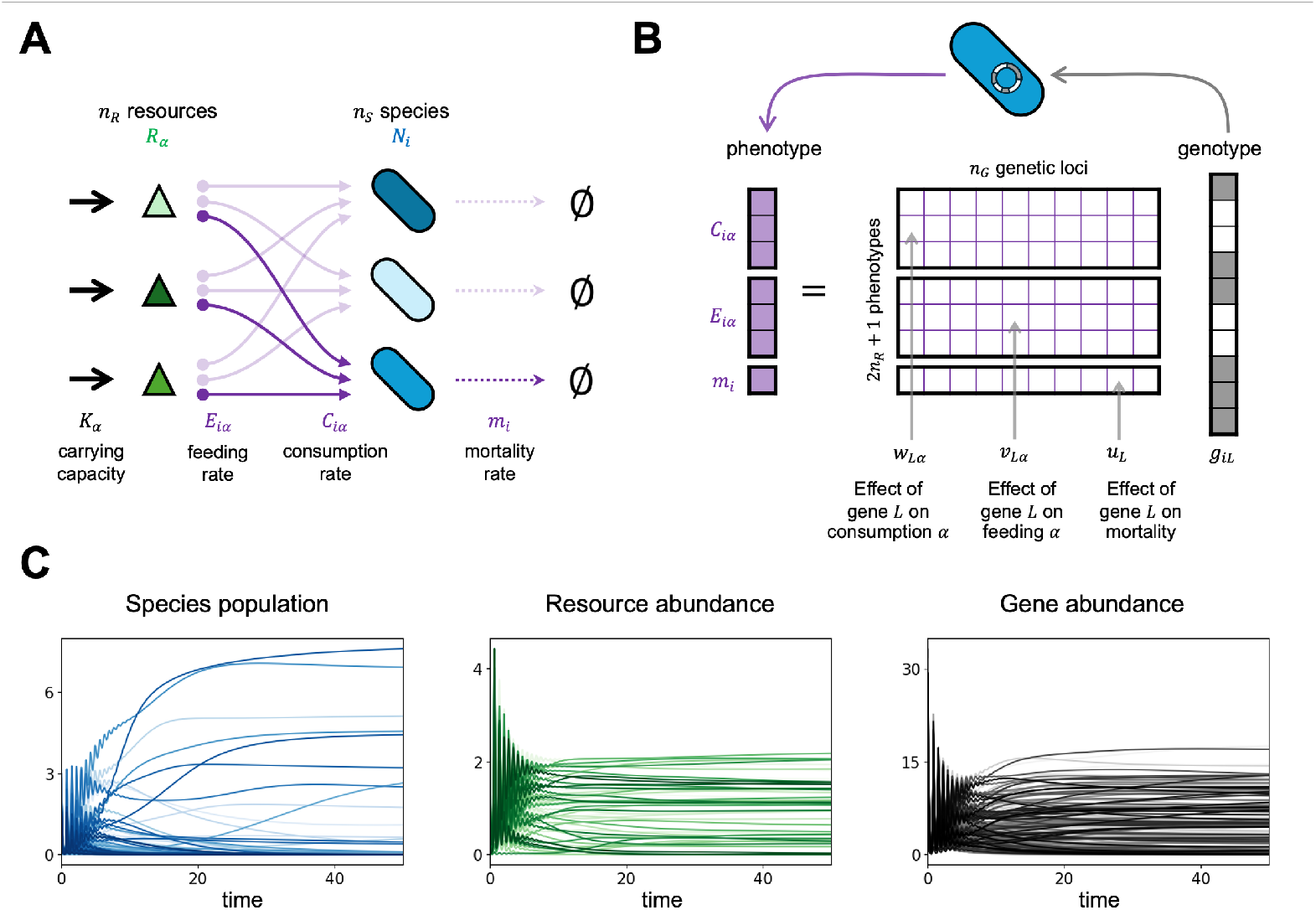
Model. (A) MacArthur’s consumer resource model. (B) Linear genotype-to-phenotype mapping. (C) Dynamics of species populations, resource abundances, and gene abundances.

- a vector of consumer preferences 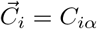 whose elements tell you the
- per-unit-resource growth rate of species *i* due to consumption of resource *α*.
- a vector of impacts 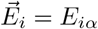 whose elements encode the per-capita depletion rate of resource *α* by species *i*. Note that in the MCRM, it is commonly assumed that *E*_*iα*_ = *C*_*iα*_, in which case the ecological dynamics have a Lyapunov function and can be formulated in terms of constrained optimization.
- a species-dependent minimum maintenance energy or death rate *m*_*i*_ that sets a lower bound on the amount resources a species must consume in order to survive in the environment.

Resources grow logistically with a resource-dependent carrying capacity *K*_*α*_ and are consumed by species that use the energy contained in the resources to grow. This gives rise to implicit competition between species, with species whose impact and preference vectors are more similar competing more strongly. This competition results in competitive exclusion between similar species, and bounds the total steady-state diversity of the ecosystem to be less than the number of resources *n*_*R*_ [17].

To incorporate G→P maps into the MCRM, we re-parameterize the phenotypic traits of each species in terms of an underlying genotype **g**. We take **g** to be an *n*_*G*_-dimensional genotype vector, with each element *g*_*L*_ of the vector corresponding to a different gene/locus. For simplicity, in our simulations we will often assume that the elements of **g** are binary (i.e. *g*_*L*_ = 0, 1), corresponding to the case where each gene has two possible alleles. However, we emphasize that our analysis holds quite generally and does not depend on this particular choice.

As in the original MCRM, each species is characterized by its phenotypic parameters, namely its consumer preferences 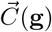, impacts 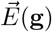, and maintenance energy *m*(**g**), now assumed to be functions of the genotype [18]. With these changes, the ecological dynamics in Eq (1) can be written as

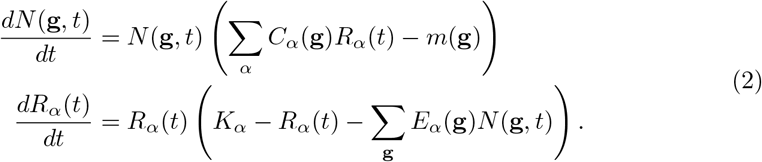

These equations make clear that the population dynamics can be viewed directly in gene space.

To make the model analytically tractable, throughout this study we assume that the phenotypic parameters are linear functions of the genotype **g**. This approximation is standard for quantitative traits governed by many loci with small, independent effects [14, 15]. Mathematically, this assumption takes the form

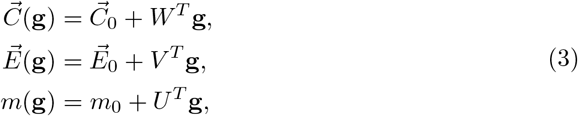

Where 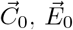, *m*_0_ represent the baseline phenotypes, and *W, V*, and *U* are the weight matrices that map genes to phenotypic traits (see Fig 1B). Each entry in these weight matrices quantifies the contribution of gene *L* to a particular phenotypic trait. Note that *W* = (*w*_*Lα*_) and *V* = (*v*_*Lα*_) are matrices of size *n*_*G*_ *× n*_*R*_, while *U* = (*u*_*L*_) is a matrix of size *n*_*G*_ *×* 1.

Fig 1C shows a typical simulation of the dynamics resulting from Eq (2). In addition to the abundances of species and resources, one can also track the abundances of individual genes in the ecosystem, a quantity that can be experimentally measured using metagenomic time-series.

### Population and resource dynamics in metagenomic terms

Given the linear G→P maps, we can analytically examine the population and resource dynamics in metagenomic terms. At different positions in the gene space, populations experience heterogeneous growth rates. We define the instantaneous growth rate of a genome as (see Fig 2A)

**Fig 2.**
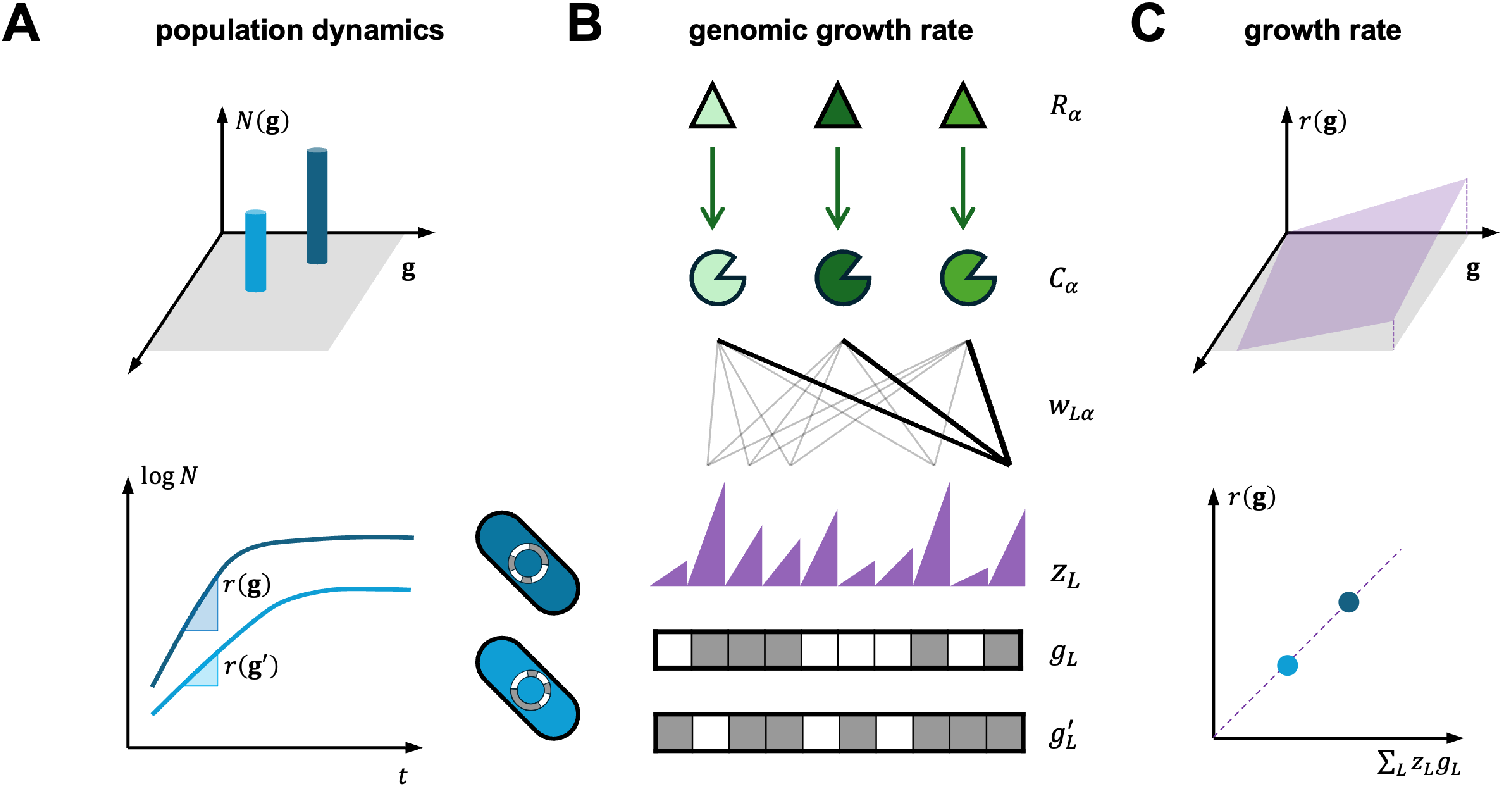
Reinterpreting ecological dynamics in terms of metagenomes. (A) Species are parameterized by their genomes **g** and have an abundance *N*(**g**) and instantaneous growth rate *r*(**g**). (B) The growth rate of gene *L* in the ecosystem, *z*_*L*_, depends on the instantaneous resource abundances *R*_*α*_ through the matrix *W*_*Lα*_ that measures how much gene *L* contributes to the consumption of resource *α* (see Eq (7)). (C) The growth rate of a species with genome **g** is the dot product of its genomic vector **g** and the vector gene-level growth rate **z**. (Eq (8)).

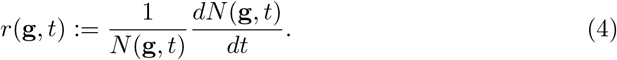

In the MCRM, this growth rate depends on the current environmental state through the resource vector 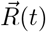:

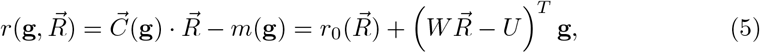

where we have defined the baseline growth rate 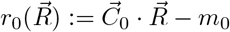. Note that 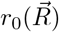 is independent of the species genotype but does depend on the current environment through the instantaneous resource abundances.

It is helpful to further decompose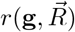 in terms of the contributions made by individual genes 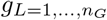 (see Fig 2B). To do so, notice that we can rewrite the second term in Eq (5) as

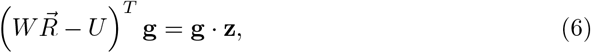

where we have defined the vector 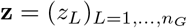 whose entries are the contribution of gene *L* to the instantaneous growth rate of a species that contains this gene:

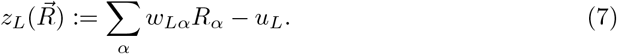

With these definitions, the instantaneous growth rate of a species with genome **g** in the ecosystem can be expressed as the baseline rate plus the collective contribution of its constituent genes:

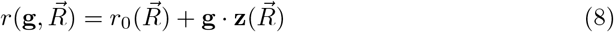

Biologically, **z** quantifies how each gene contributes to fitness under the current resource composition. It represents the ecological selective pressure acting on each gene within the metagenomic landscape.

Using the method of integrating factor, the population dynamics can then be solved explicitly as

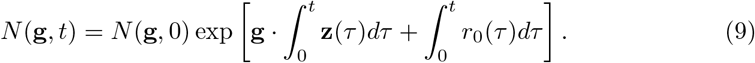

This solution describes how the population distribution evolves and shifts across the gene space, driven by the cumulative ecological selective forces encoded in 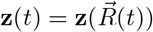.

As for the resource dynamics, each resource follows logistic growth with an effective carrying capacity that evolves over time as the community composition changes. Specifically,

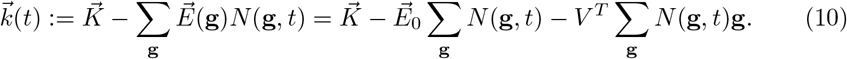

We see that the temporal carrying capacity depends linearly on two statistical quantities describing the community populatLion: (a) the total biomass *B*(*t*):= ∑_**g**_ *N*(**g**, *t*); and (b) the gene abundances *G*_*L*_(*t*):= ∑_**g**_ *N*(**g**, *t*)*g*_*L*_. It is convenient to introduce the mean gene abundance vector of the community, 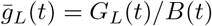, which represents the average genetic composition weighted by population size. Substituting this into the expression for 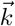, the effective carrying capacity can be rewritten as

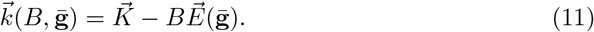

This form shows that the available resources are jointly regulated by the total biomass and the average genomic traits of the community. Finally, the resource dynamics can be explicitly integrated as

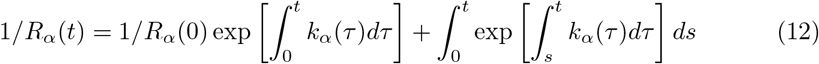

Together with Eq (9), these results show that population dynamics and resource dynamics are tightly coupled: the resource availability shapes growth rates through the metagenomic selective force 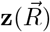, while the evolving community metagenomic statistical quantities in turn modulates the resource environment through the effective resource carrying capacity 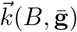.

### Dynamics of metagenomic statistics

To understand how ecological and genomic processes are coupled, we next examine the dynamics of the statistical quantities that describe the metagenome. These quantities summarize the population distribution in gene space and capture how the overall genomic composition of the community evolves under ecological feedback. To this end, we introduce the moment-generating function (MGF) of the population distribution,

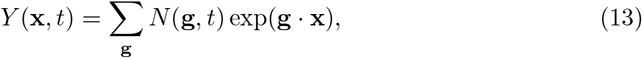

and its logarithm, the cumulant-generating function (CGF),

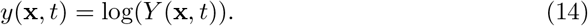

These functions compactly encode all statistical moments and cumulants of the community metagenome. The total biomass of the community corresponds to the MGF evaluated at the origin, *B*(*t*) = *Y* (**x**, *t*)|_**x**=**0**_. The abundance of gene *L* is given by the first moment, 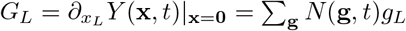, while the mean genome of the community is given by the first cumulant 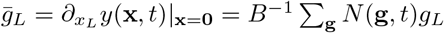. Higher-order derivatives of these functions allow us to derive statistics about how often genes co-occur in the same species. For example,

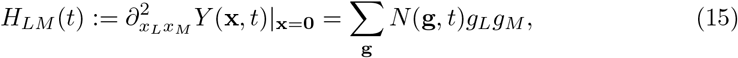

which quantifies how often gene *L* and gene *M* co-occur within the same genomes in the community.From the population growth equation in Eq (2), we can derive the governing equations for MGF and CGF:

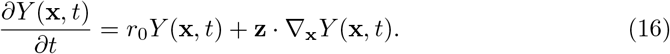

and

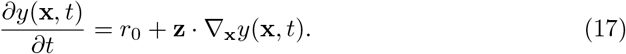

Eq (16) and Eq (17) describe how the statistical moments and cumulants of the metagenome evolve over time under the influence of resource-dependent selection gradients.

Evaluating Eq (16) at **x** = 0 gives the dynamics of the total biomass

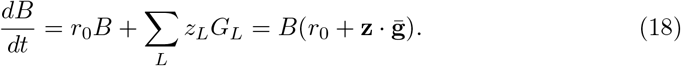

This equation shows that the total biomass grows at a rate determined by the growth rate evaluated at the mean genome 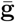. Similarly, the dynamics of the gene abundances are obtained by differentiating Eq (16) with respect to *x*_*L*_, leading to

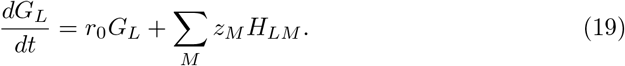

To characterize correlations beyond mean abundances, we define the covariance (second cumulant) of the metagenome as

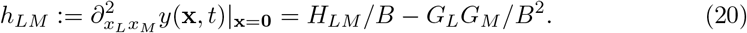

This covariance, which we refer to as metagenomic hitchhiking, measures the extent to which the presence of gene *L* predicts the presence of gene *M* beyond what is expected by chance. From Eq (17), the dynamics of the mean genome are then given by

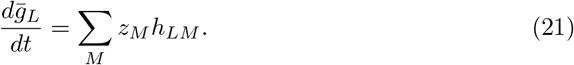

This expression shows that the mean genome evolves along directions in gene space where the gene–gene covariance structure aligns with the environmental selection gradient **z**. In other words, selection drives shifts in the community’s average genomic composition through correlated gene dynamics.

Together, Eq (18) and Eq (21) reveal a hierarchical structure of metagenomic dynamics: the growth of total biomass depends on the mean genome, the mean genome evolves through gene covariances, and these covariances are shaped by the underlying population distribution in gene space. This hierarchical coupling provides a compact and interpretable description of how ecological feedback drives the evolution of metagenomic statistics.

### Metagenomic fitness and competition in the effective GLV system

The feedback between organisms and the environment gives rise to a coupling between community composition and metagenomic features. To better understand how metagenomic traits influence community-level selection and competition, it is useful to examine the limit where environmental variables equilibrate rapidly relative to population changes. In this quasi-steady-state regime, resource dynamics satisfy *dR*_*α*_*/dt* = 0, and all resources persist with abundances determined by the effective carrying capacity in Eq (11).

With these substitutions, the species dynamics take the form of a Generalized Lotka–Volterra (GLV) system with species labeled by genomes **g**:

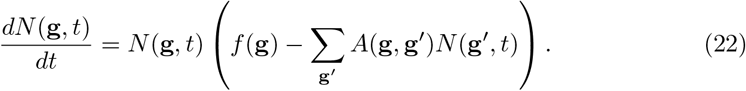

For notational simplicity, we restrict ourselves to the case where 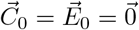 and *m*_0_ = 0 in Eq (3), in which case, the species fitness takes the form

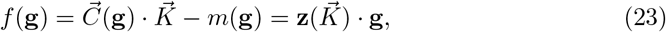

and the competition kernel between species **g** and **g**^*′*^ is given by

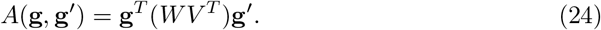

Notice that the fitness *f*(**g**) of an species with genome **g** is a linear sum of contributions from each gene

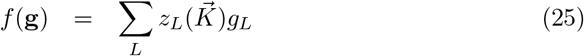

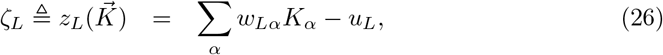

allowing us to assign a gene-level fitness *ζ*_*L*_ to each gene. Similarly, we can write the competition kernel between species *A*(**g, g**^*′*^) as a sum of gene-level competition kernels as

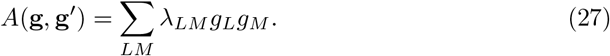

with

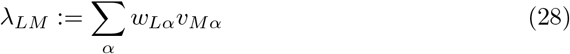

the effective gene-gene interaction matrix.

In this effective GLV system, the dynamics of the total biomass take the simple form

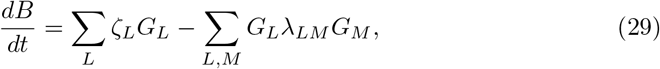

which is governed by the current gene abundances *G*_*L*_. If the system is reciprocal, i.e., *W* = *V*, the competition matrix *λ*_*LM*_ is symmetric. In this case, the dynamics admit a Lyapunov function

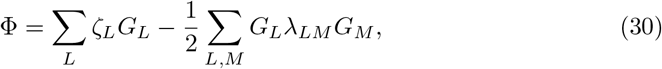

which guarantees convergence to a stable equilibrium. The system evolves toward maximizing the aggregate metagenomic fitness while minimizing total competitive load, reflecting a balance between selection for beneficial genes and constraints from shared resource utilization.

This formulation highlights how ecological interactions and genomic composition are interlinked: community dynamics can be interpreted as the gradient ascent of a metagenomic fitness landscape. The equilibrium reflects a balance between the selective amplification of beneficial genes and the suppression imposed by shared competition. Hence, even in a static environment, the structure of the metagenome dynamically encodes the outcome of eco-evolutionary feedback between selection and resource-mediated constraints for linear G→P maps.

## Results

To better understand how ecology shapes the metagenomic properties of ecosystems, we carefully analyzed the MacArthur consumer-resource model (MCRM) in Eq (2). We restricted ourselves to the case where the interactions are reciprocal (*C* = *E*), and the ecological dynamics reach a unique steady-state. We parameterized genomes **g** as binary vectors, 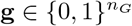, indicating the presence or absence of a gene.

To investigate the typical properties of large ecosystems, we focused on the case where the parameters *w*_*Lα*_ and *u*_*L*_ in Eq (3) defining the G→P map were drawn randomly, i.e., 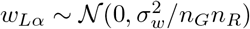 and 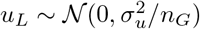. For simplicity, we assumed that the environment was homogeneous with all *K*_*α*_ = *µ*_*K*_ and the baseline parameters 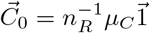 all fixed constants.

Through these simulations, we analyzed how metagenomic properties shape community organization. First, we examined how gene fitness and metagenomic hitchhiking jointly determine gene survival. Next, we explored species coexistence in communities whose genomes exhibit hierarchical structure, revealing how genetic correlations affect diversity. Finally, we investigated the ecological steady state in systems with low-rank genotype-to-phenotype maps, corresponding to coarse-grained biochemical pathway organization.

### Gene survival determined by gene fitness and metagenomic hitchhiking

We began by examining which genes survived or went extinct once the ecological dynamics converged to a steady state. Under the random parameterization described above, the gene-level fitness values *ζ*_*L*_ (Eq (28)) were approximately Gaussian-distributed with zero mean. In a randomly assembled community with *n*_*S*_ species, each gene appeared in any given genome with probability *p*_*g*_. Across simulations of many such communities (see Methods), we observed that the mean gene abundance ⟨*G*_*L*_⟩ increased with gene fitness: high-fitness genes consistently reached higher abundances at steady state (Fig 3A). Likewise, the probability of gene survival, *P*(*G*_*L*_ *>* 0), was positively correlated with *ζ*_*L*_ (Fig 3B), consistent with standard ecological intuition.

**Fig 3.**
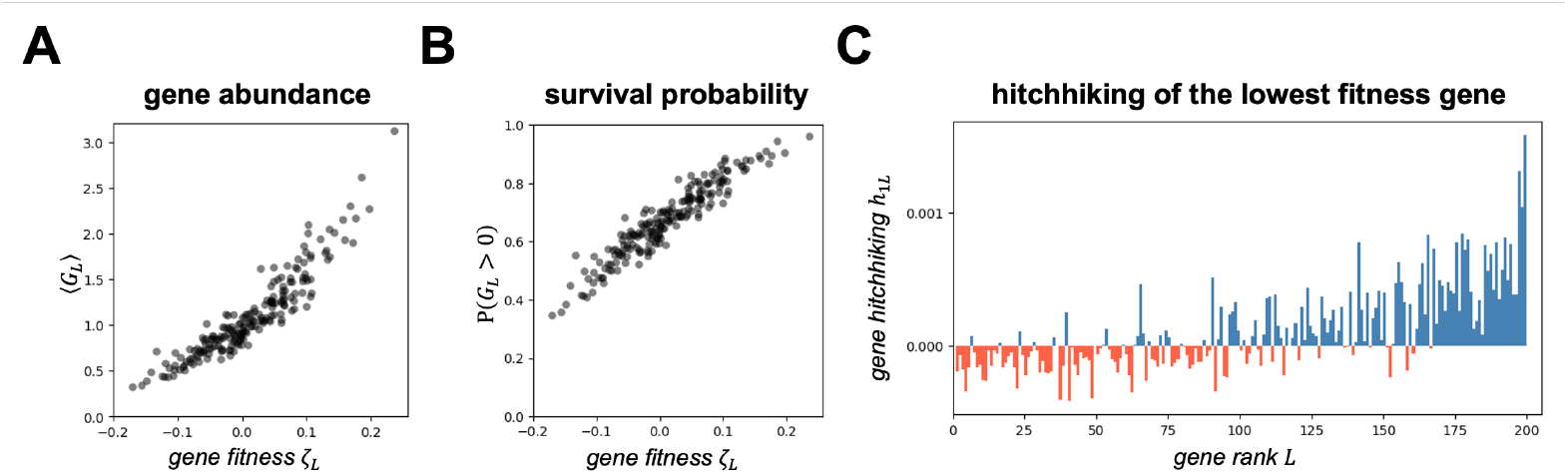
Gene survival determined by gene fitness and metagenomic hitchhiking. (A) The average gene abundances ⟨*G*_*L*_⟩ correlate with gene fitness *ζ*_*L*_. (B) The probability of a gene to survive *P*(*G*_*L*_ *>* 0) correlates with gene fitness *ζ*_*L*_. (C) The lowest-fitness gene (*L* = 1) co-occurs with high-fitness genes above expectation when it survives. Gene indices *L* are ranked from low to high fitness.

However, a key difference emerged between gene-level and species-level selection. Even the lowest-fitness genes retained a survival probability above 20%. This contrasted with species dynamics, where species with sufficiently low fitness reliably went extinct when competitive interactions were approximately homogeneous. This discrepancy arose because gene fitness acted through the genomes of species: a gene with intrinsically low fitness could survive if it appeared in species whose overall fitness was high. In other words, low-fitness genes could persist through metagenomic hitchhiking, in analogy with the “hitch-hiking effect” in population genetics [19].

To demonstrate this mechanism, we examined the second cumulant of the metagenome (Eq (20)), which quantifies gene–gene co-occurrence above expectation. As shown in Fig 3C, whenever the lowest-fitness genes survived, they displayed systematically stronger hitchhiking with high-fitness genes than with other low-fitness genes. Thus, survival was driven not only by intrinsic gene fitness but also by the fitness of the genomic backgrounds in which genes resided.

In summary, a gene survived either because it was intrinsically fit or because it was carried by fit species. This metagenomic persistence — enabled by genomic context — is a fundamental feature of ecological dynamics, allowing otherwise unfit genes to be maintained in the community and potentially become beneficial under future environmental change.

### Species coexistence in communities with phylogenetic tree structure

Incorporating G→P maps into ecological models enabled us to study communities whose species had structured genomes, particularly those reflecting phylogenetic relationships. In such settings, closely related species share more genes due to common ancestry, and hence exhibit more similar phenotypes (e.g., resource preferences). To examine how phylogenetic structure shaped coexistence, we generated communities in which species genomes arose by iterated mutation along prescribed tree topologies (see Methods). We analyzed three representative cases: a perfectly unbalanced binary tree (Fig 4A), a perfectly balanced binary tree (Fig 4D), and a multi-family balanced tree composed of several independent balanced clades (Fig 4G).

**Fig 4.**
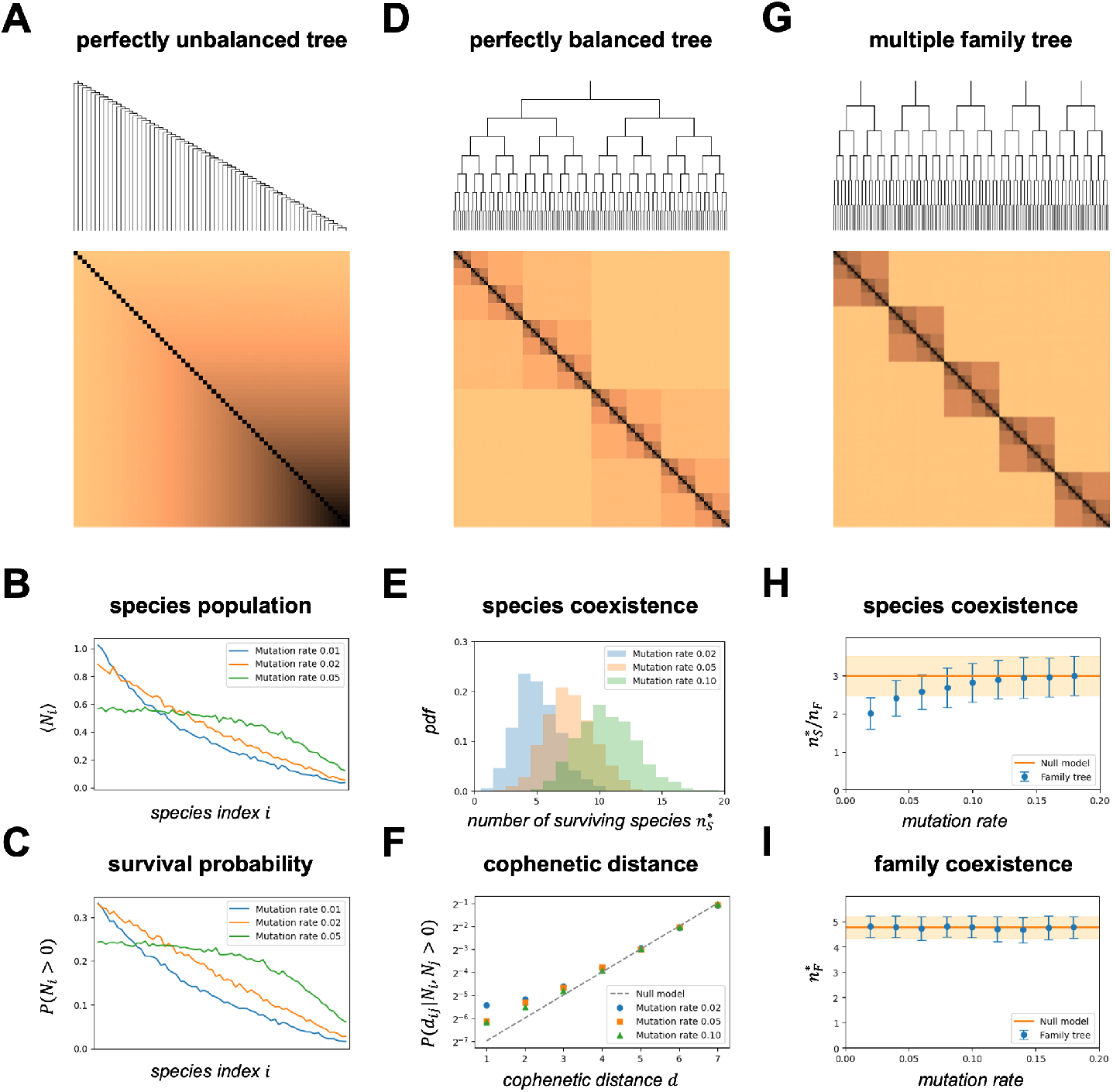
Species coexistence in communities with phylogenetic tree structures. (A) The phylogenetic graph of a perfectly unbalanced tree and the genomic distances between species. (B) The left species in the tree has higher populations than the right species. (C) The left species in the tree has higher probabilities of surviving than the right species. (D) The phylogenetic graph of a perfectly balanced tree and the genomic distances between species. (E) Higher mutation rates enable more coexisting species. (F) The probability distribution of the cophenetic distance between coexisting species. (G) The phylogenetic graph of a multiple-family balanced tree and the genomic distances between species. (H) The number of coexisting species per family. (I) The number of coexisting families. Orange lines and shades are the mean and the standard deviation of the null model.

#### Perfectly unbalanced tree

In the unbalanced tree, a pool of *n*_*S*_ species was generated through *n*_*S*_ − 1 consecutive asymmetric branching events, each occurring along the rightmost lineage. The leftmost species thus sat deepest in the tree and was maximally distant (in genomic space) from all others, whereas species became progressively more similar moving toward the right. Because genomic similarity induces stronger overlap in resource use, closely related species experienced stronger competitive interactions. Simulations of ecological dynamics revealed a clear pattern: the leftmost (most distinct) species had the highest probability of surviving, while species near the rightmost tip were the most extinction-prone (Fig 4B). Moreover, the expected steady-state population sizes declined monotonically from left to right (Fig 4C). Both findings were consistent with classical predictions that closely related species compete more intensely and therefore have reduced equilibrium abundances. Increasing the mutation magnitude at branching events weakened competition, which increased both survival probabilities and steady-state population sizes, especially for the compact side of the tree.

#### Perfectly balanced tree

In the balanced tree, *b* symmetric branching events produced *n*_*S*_ = 2^*b*^ species, each with identical depth and identical distribution of genomic distances to others. As expected from symmetry, all species exhibited identical survival probabilities and identical expected equilibrium abundances (Fig 4E). These quantities again increased with mutation rate, reflecting reduced niche overlap. We also examined the cophenetic distance — defined as the number of branching steps required for two species to meet at a common ancestor — between surviving species. Under a null model where survivors were chosen uniformly at random, the probability that two survivors had cophenetic distance *d* equaled 2^*d*−*b*^. Intuitively, one might expect competitive exclusion to favor coexistence of phylogenetically distant species, leading to *P*(*d*) *>* 2^*d*−*b*^ for large *d*. Surprisingly, the simulations showed the opposite trend: surviving species were more likely than random to be close relatives, including sister species (Fig 4F). A plausible explanation is that branching events often produced daughter species with systematically different fitness, biasing survival toward the higher-fitness branch and thereby reducing cophenetic distances among survivors.

#### Multiple-family balanced tree

In the multi-family scenario, the community consisted of *f* phylogenetically isolated families, whose ancestral genomes shared no overlap. Within each family, species arose from *b* balanced branching steps, yielding *n*_*S*_ = *f ×* 2^*b*^ species. Because inter-family genomic distances are large, inter-family competition was weak, allowing multiple families to coexist. Within each family, multiple species could coexist as well, and the number of within-family survivors increased with mutation rate (Fig 4H), again reflecting reduced competition. Notably, however, the number of surviving families remained essentially insensitive to mutation rate (Fig 4I), suggesting a form of hierarchical ecological packing: family-level coexistence was determined primarily by coarse-grained genomic structure, whereas species-level coexistence within families depended more sensitively on mutation-induced variation.

In summary, these results showed that incorporating G→P maps into ecological dynamics provided a principled framework for understanding how phylogenetic relationships shape community composition. Phylogenetic topology, mutation magnitude, and genomic distance together determined both the number and identity of surviving species. This framework naturally connects ecological coexistence with evolutionary history, highlighting how lineage structure and competitive interactions jointly organize eco-evolutionary communities [20].

### Ecological steady state for low-rank G→P maps

In biological systems, phenotypes are often structured by a limited number of functional pathways. Multiple resources may be processed through the same metabolic module, and a single gene can influence multiple traits via shared regulatory networks. As a result, realistic genotype-to-phenotype (G→P) maps are expected to be low-rank, reflecting the fact that cellular processes are governed by a relatively small number of effective biochemical pathways [21, 22].

Our framework naturally allows us to model this situation by imposing a low-rank structure on the weight matrices *W*. Starting from the Gaussian random matrices described above, we constructed low-rank G→P maps by truncating small singular values in their singular value decomposition. The resulting rank *n*_*P*_ can be interpreted as the number of effective pathways through which genomes modulate resource consumption and environmental impact.

Simulations revealed a clear constraint on community diversity: the number of coexisting species at ecological steady state could not exceed the phenotypic rank, achieving at most *n*_*P*_ + 1 survivors. This behavior generalizes the classical competitive exclusion principle, indicating that coexistence is bounded not by the number of resources per se, but by the dimensionality of the phenotypic space induced by the G→P map. As the number of pathways *n*_*P*_ increased, the observed diversity began to deviate from the phenotypic rank bound (Fig 5B). This deviation occurred later when the environment was richer — i.e., when the carrying capacities *K*_*α*_ were large — because high resource availability reduces effective competition among pathways, delaying saturation of the coexistence limit. Conversely, increasing *n*_*P*_ also led to a higher number of resource extinctions at steady state (Fig 5C), reflecting that a greater diversity of phenotypes drives stronger aggregate pressure on the resource pool.

**Fig 5.**
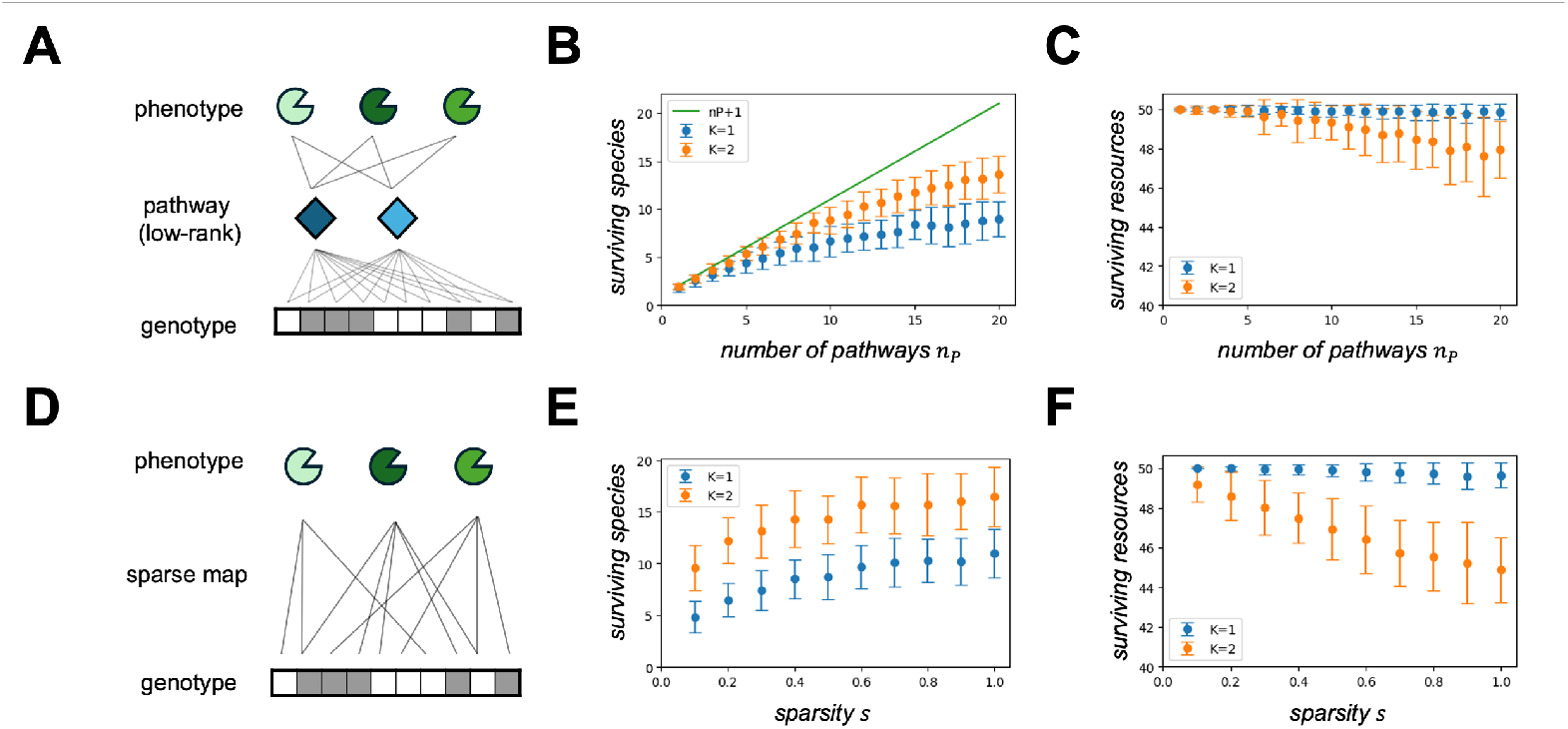
Ecological steady state for sparse or low-rank G→P maps. (A) Low rank genotype-to-phenotype maps through pathways. (B) Number of surviving species under low-rank genotype-to-phenotype maps with different resource carrying capacities. (C) Number of surviving resources under low-rank genotype-to-phenotype maps with different resource carrying capacities. (D) Sparse genotype-to-phenotype maps. (E) Number of surviving species under sparse genotype-to-phenotype maps with different resource carrying capacities. (F) Number of surviving resources under sparse genotype-to-phenotype maps with different resource carrying capacities.

### Ecological steady state for sparse G→P maps

According to the biophysical nature of cells, most genes are specialized for only a few functions. As a result, realistic G→P maps are expected to be sparse, meaning that each gene has non-zero contributions to only a small number of phenotypes. In our framework, we introduce the sparsity of the G→P map, denoted by *s*, to represent the proportion of non-zero weights in the matrix *W*, as illustrated in Fig. 5D. Intuitively, a sparse G→P map implies that different species tend to occupy more similar positions in phenotype space, thereby reducing phenotypic diversity. Simulations confirm this theoretical intuition. Fig. 5E shows that fewer species can coexist when the G→P map is more sparse, a trend that holds for both high resource supply (*K* = 2) and low resource supply (*K* = 1). Fig. 5F shows that more resources can coexist under sparse G→P maps, with the trend being particularly clear when the resource supply is high (*K* = 2).

## Discussion

By incorporating linear genotype-to-phenotype maps into ecological modeling frameworks, we reformulated community dynamics in terms of metagenomic variables. We showed that effective resource carrying capacities are determined by the total biomass and the mean genome of the community, and that population dynamics are governed by the alignment between species genomes and metagenomic selection forces. Furthermore, the evolution of total biomass and mean genome is influenced by higher-order metagenomic cumulants, such as gene hitchhiking, and ecological steady states are shaped by fitness and competition defined at the genomic level.

Through numerical simulations, we demonstrated that genes with high intrinsic fitness had a higher probability of persistence, while low-fitness genes could nonetheless survive by hitchhiking within high-fitness genomes. The framework further enabled systematic investigation of ecological patterns observed in microbial systems, including communities structured by phylogenetic trees and systems characterized by low-rank or sparse G→P maps. Together, these results highlight the broad applicability of our approach for linking genomic architecture to ecological organization.

### Theoretical implications

Our mathematical framework reveals rich dynamical structures that couple genomic and species-level processes, opening several directions for future theoretical work. By explicitly resolving genotypes and phenotypes, the model emphasizes that a cell is not a simple input–output unit but a composite system comprising multiple multifunctional genes. This perspective naturally captures trade-offs, correlations, and coexistence mechanisms that are central to community ecology, while also giving rise to a hierarchy of metagenomic moments that may be of independent interest from a statistical physics standpoint [23–25].

More broadly, the framework raises questions concerning the emergence of species and strains, as well as the eco-evolutionary processes that shape them [26]. Ecological and evolutionary dynamics are jointly driven by mutation and horizontal gene transfer, both of which can be naturally incorporated into our formulation [27, 28]. While the present work focuses on linear G→P maps, extending the model to include quadratic or more general nonlinear mappings — thereby capturing gene–gene interactions, epistasis, or cooperative effects — represents an important direction for future research [29, 30].

### Experimental implications

Our study provides a bridge between ecological theory and experimental biology by focusing on metagenomic quantities that are directly accessible in microbial systems. For example, measurements of gene-level growth rates in metagenomic data could be used to infer metagenomic selection forces acting within a community, thereby providing indirect information about environmental constraints and resource availability [25, 31–33]. Such inference could, in turn, enable predictions of coexistence and steady-state composition in newly assembled or perturbed communities.

Conversely, by analyzing steady-state community compositions across multiple environments, one may infer gene fitness values and effective competition between genes, offering a potential route to predicting latent or unknown gene functions [34]. This framework also underscores the importance of sequencing technologies that preserve linkage information among genes, which are essential for quantifying metagenomic hitchhiking effects. Although the model necessarily simplifies the complexity of real microbial ecosystems, integration with experimental data will allow parameter calibration and refinement, ultimately enhancing the predictive power and biological relevance of the theory.

## Methods

All simulations were implemented in *Jupyter Notebook*. The code is available on GitHub: https://github.com/Emergent-Behaviors-in-Biology/Ecology_of_metagenomes.

### Parameter setup

Given the number of resources *n*_*R*_ and genes *n*_*G*_, we sample the genotype-to-phenotype weight matrix *W* = (*w*_*Lα*_) of size *n*_*G*_ *× n*_*R*_ from a Gaussian distribution

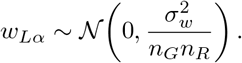

For the reciprocal MCRM, we set *V* = *W*. In addition, the weight vector *U* = (*u*_*L*_) of size *n*_*G*_ *×* 1 is sampled from the Gaussian distribution

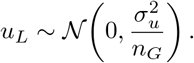

Given the number of species *n*_*S*_, the genome of species *i*, **g**_*i*_ = (*g*_*iL*_), is sampled from a Bernoulli distribution

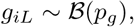

with probability *p*_*g*_ for gene presence. The parameters *µ*_*C*_ and *µ*_*K*_ determine the mean values of consumer preference and resource supply, respectively.

Unless otherwise stated, simulations use the hyperparameters listed in Table 1. Some parameters are varied in later sections.

**Table 1.** The default value of hyperparameters in the simulations.

| Hyperparameter | Defination | Default value |
| --- | --- | --- |
| $n_G$ | the number of gene loci | 200 |
| $n_S$ | the number of species | 100 |
| $n_R$ | the number of resource types | 50 |
| $p_g$ | the probability of a gene present $g_L = 1$ in a genome $\mathbf{g}$ | 0.1 |
| $\sigma_w$ | the normalized standard deviation of the random matrix $W$ | 1 |
| $\sigma_u$ | the normalized standard deviation of the random matrix $U$ | 0.1 |
| $\mu_K$ | the carrying capacity of resources | 1 |
| $\mu_C$ | the baseline consumption rate | 2 |
| $m_0$ | the baseline mortality rate | 1 |

### Dynamics and equilibrium

The initial population abundances *N*_*i*_(0) and resource levels *R*_*α*_(0) are independently sampled from a uniform distribution on (0, 1). The ordinary differential equations of the MCRM are solved using the *RK45* method with time step Δ*t* = 0.01. Simulations shown in Fig. 1 are run until *t* = 50.

For the reciprocal MCRM, the equilibrium state can be obtained by solving the corresponding constrained quadratic optimization problem [35, 36]. The numerical tolerance is set to 10^−7^.

### Metagenome statistics

We perform 10,000 realizations of the MCRM equilibrium. In these realizations, the weight matrices *W* and *U* are fixed, while the species genomes *g*_*iL*_ are resampled each time. For convenience, gene indices *L* are reordered according to the rank of gene fitness, from low to high.

After all simulations, we compute the average gene abundance

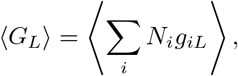

and the average gene co-occurrence

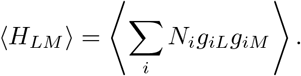

The gene hitchhiking coefficient *h*_*LM*_ is then calculated from ⟨*G*_*L*_ ⟩ and ⟨*H*_*LM*_⟩ according to Eq (20). In addition, the survival probability of gene *L* is defined as the fraction of realizations in which *G*_*L*_ *>* 0. The results are shown in Fig. 3.

### Community with unbalanced tree structure

To generate genomes with a perfectly unbalanced phylogenetic tree, we first create an ancestral genome where each gene is present with probability *p*_*g*_. At each branching step, we copy the genome of the species with the largest index to create a new species, and then mutate all genomes in the community with mutation rate *γ* ∈ (0, 1).

During mutation, each present gene (*g*_*iL*_ = 1) changes to 0 with probability *γ*, while each absent gene (*g*_*iL*_ = 0) changes to 1 with probability *γp*_*g*_*/*(1 − *p*_*g*_). This procedure preserves the expected number of present genes. After *n*_*S*_ − 1 branching events, the community contains *n*_*S*_ species following the unbalanced tree structure shown in Fig. 4A. The statistics in Fig. 4BC are obtained from 10,000 realizations.

### Community with balanced tree structure

To generate a perfectly balanced tree, we start with a single ancestral genome with gene presence probability *p*_*g*_. At each generation, all genomes in the community are duplicated and mutated with mutation rate *γ*. After *b* branching generations, the community contains *n*_*S*_ = 2^*b*^ species forming the balanced tree structure shown in Fig. 4D. The statistics in Fig. 4EF are computed from 10,000 realizations.

### Community with multiple family trees

To construct communities with multiple phylogenetic families, we begin with *n*_*F*_ ancestral genomes. These ancestors share no common present genes. For each family, a balanced tree is generated with mutation rate *γ*. After *b* branching generations, the community contains

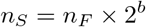

species, as illustrated in Fig. 4G. The statistics in Fig. 4HI are computed from 10,000 realizations.

### Low-rank G→P maps

To generate low-rank genotype-to-phenotype maps, we set the number of pathways (rank) to *n*_*P*_ *< n*_*R*_. The weight matrix can be written as the product *W* = *W*_1_*W*_2_, where *W*_1_ has size *n*_*G*_ *× n*_*P*_ and *W*_2_ has size *n*_*P*_ *× n*_*R*_.

To preserve the Gaussian statistics of the weights, we first generate a Gaussian random matrix *W* as described above and then perform singular value decomposition. Only the *n*_*P*_ largest singular values are retained, while the remaining singular values are set to zero. This produces a matrix *W* of rank *n*_*P*_, corresponding to a low-rank G→P map (Fig. 5A). The statistics in Fig. 5BC are computed from 100 realizations for each parameter set.

### Sparse G→P maps

To generate sparse G→P maps, we introduce a sparsity parameter *s*, defined as the fraction of non-zero entries in the weight matrix *W*. After generating a Gaussian random matrix *W*, each entry is independently set to zero with probability 1 − *s*. This procedure produces sparse weight matrices as shown in Fig. 5D. The statistics in Fig. 5EF are computed from 100 realizations for each parameter set.

## Acknowledgments

We would like to thank members Zhijie Feng, Akshit Goyal, Kobe Li, and members of the Mehta group for useful discussions. This work was supported by NIH NIGMS R35GM119461 and a Chan-Zuckerberg Investigator grant to PM.

## Notes

https://github.com/Emergent-Behaviors-in-Biology/Ecology_of_metagenomes.

### Competing Interest Statement

The authors have declared no competing interest.

